# Recessive *POPDC1* Truncation Causes Lethal Short-QT Pattern Arrhythmogenic Cardiomyopathy with Multi-Ion Channel Remodeling and Ankyrin-G Scaffold Disruption

**DOI:** 10.64898/2026.03.07.710328

**Authors:** Rong Luo, Chenqing Zheng, Huan Lan, Yilin He, Yi Wang, Qin Sheng, Shuang Li, Hongmei Deng, Lei Yao, Yifei Li, Wei-Wen Lim, Wei Hua, Xiushan Wu, Xin Cao, Xiaoping Li

## Abstract

**Aims:** Biallelic variants in *Popeye domain containing 1* (*POPDC1*) classically cause limb-girdle muscular dystrophy, but their impact on cardiac system remains unclear. We investigated the functional consequence of a POPDC1 frameshift variant (c.448delT), first identified in a consanguineous family with arrhythmogenic cardiomyopathy (ACM) and sudden death.

**Methods and results:** Comprehensive clinical and genetic evaluation was followed by mechanistic studies in an orthologous *Popdc1* knock-in rat model. Functional characterization included biotelemetry and programed electrical stimulation, optical mapping, patch clamp, intracellular Ca^2+^ imaging, proteomics, and oxidative stress assays. Homozygous mutants (*Popdc1*^fs/fs^) exhibited a shortened QTc interval and high incidence of ventricular tachycardia compared to wild-type (*Popdc1*^+/+^). Urethane anesthesia provoked second-degree AV block in all 10 *Popdc1*^fs/fs^ rats but in only 1 of 9 *Popdc1*^+/+^ littermates. Optical mapping demonstrated abbreviated action potentials, slowed conduction velocity, and inducible polymorphic VT. Patch clamp confirmed accelerated repolarization, with upregulated transient outward potassium currents (*I*_to_) and L-type calcium currents (*I*_Ca, L_), but downregulated peak sodium currents (*I*_Na_). Multi-omics and ultrastructural analyses revealed a post-translational collapse of intercalated disc: POPDC1 loss destabilized Ankyrin-G and its membrane-anchored cargo, Nav1.5, coinciding with enhanced Kv4.3 and Cav1.2 protein levels. These disruptions create a convergence of delayed conduction and shortened refractoriness, forming substrate for malignant re-entry.

**Conclusion:** We defined a recessive short-QT ACM leading to potentially fatal arrhythmias caused by biallelic POPDC1 truncation driving Ankyrin-G disruption, manifesting as a triad: (1) bradycardia/AV block, (2) accelerated repolarization as short-QT pattern/diffuse T-wave flattening or inversion, and (3) progressive cardiomyopathy and sudden death.

**What’s new?:**

1. We define a novel diagnostic triad for recessive short-QT arrhythmogenic cardiomyopathy (rSQT-ACM) in a loss-of-function *POPDC1* truncation: (1) early-onset bradyarrhythmia (sinus bradycardia or AV block); (2) a paradoxical short-QT interval with maladaptive rate response/diffuse T-wave flattening or inversion; and (3) progressive cardiomyopathy accompanied accompanied by with SCD.
2. Mechanistically, biallelic *POPDC1* truncation induces collapse of the POPDC1-AnkG hub, which triggers a unique “opposing remodeling” phenotype (severe *I*_Na_ reduction coupled with paradoxical upregulation of Kv4.3-mediated *I*_to_), and established a unified “slow-conduction, short-wavelength” substrate that underpins malignant re-entry.
3. Implications for safety and device therapy: Nav1.5 deficiency indicates that sodium channel blockers may be contraindicated, and early ICD implantation may be preferable to bradycardia pacing.
4. Mechanism-directed treatments: POPDC1’s recessive inheritance and small gene size make it a strong candidate for AAV-mediated gene supplementation aimed at restoring the intercalated disc architecture and ion-channel anchoring.

Graphical Abstract

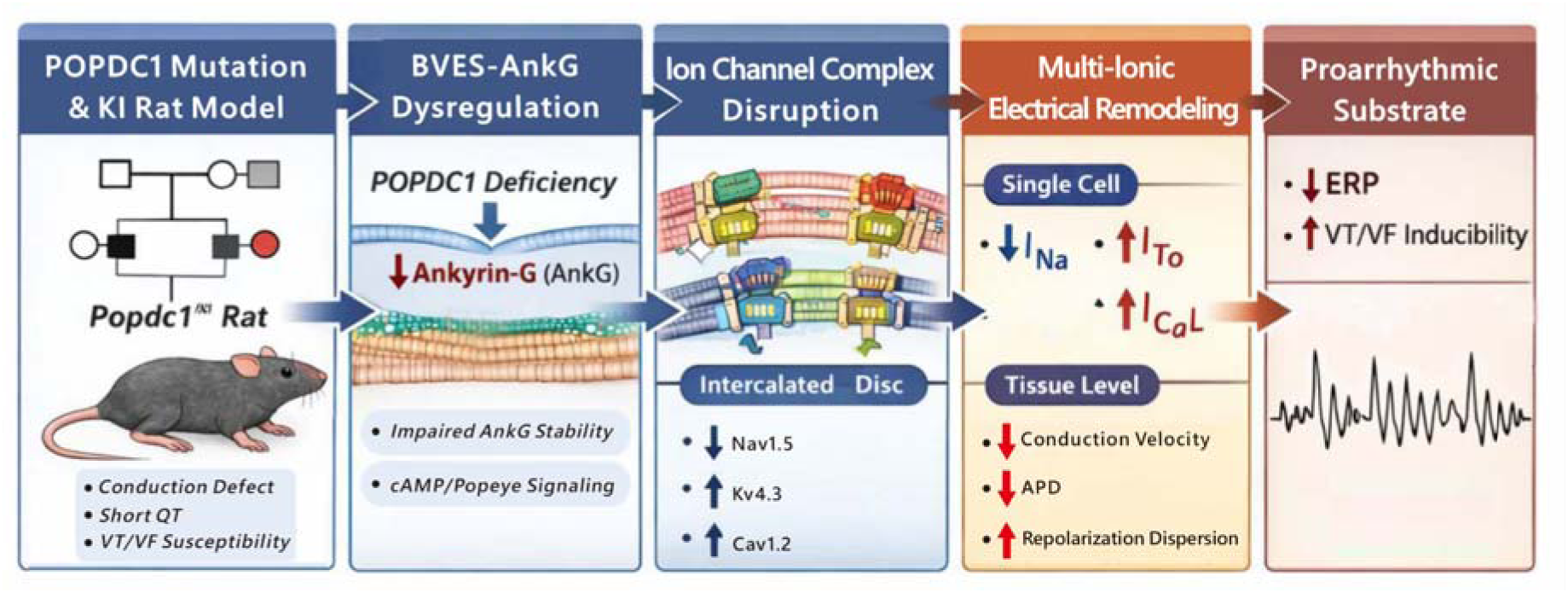

## Introduction

Arrhythmogenic cardiomyopathy (ACM) and sudden cardiac death (SCD) remain leading causes of mortality, often arising from perturbations in cardiac electrophysiology impairing electrical impulse initiation and propagation. Normal rhythm requires precise ion-channel activity and cardiomyocyte macromolecular complexes’ structural integrity, which collectively determine action potential generation, conduction velocity, and refractory dynamics (1). It is well-known that genetic variants disrupting these processes can predispose individuals to malignant arrhythmias despite overt structural heart disease.

The Popeye domain-containing (POPDC) family includes POPDC1 (also known as BVES), POPDC2, and POPDC3. *POPDC* encodes transmembrane proteins predominantly enriched in striated muscle and the cardiac conduction system (1). These proteins function as cAMP effector proteins and scaffolds that stabilize macromolecular complexes involved in electrical signaling (2, 3). Loss-of-function mutations in *Popdc* genes in animals cause heart rate variability, conduction disturbances, and stress-induced bradyarrhythmias, underscoring their importance in pacemaking and conduction system stability (1, 2, 4). Despite the classical link to limb-girdle muscular dystrophy (LGMDR25) (4, 5), emerging clinical evidence shows that POPDC1 loss can produce profound cardiac electrical abnormalities, including atrioventricular (AV) block and bradycardia (4, 6).

Despite these insights, the mechanisms by which POPDC1 dysfunction drives malignant arrhythmogenic phenotypes remain unclear. The POPDC1 complex, comprising multiple cardiac proteins, maintains electrical and structural nanodomains critical for action potential propagation (1, 7). As a chaperone, it recruits partners Caveolin-3 and two-pore potassium channels (TREK-1) to regulate vesicular trafficking and membrane retention of ion channel complexes (2,8), POPDC1 depletion associates with maladaptive conduction remodeling (9,10,11), creating a structural milieu conducive to re-entrant arrhythmias. Crucially, *Popdc1* loss-of-function in zebrafish report species-specific prolonged repolarization and calcium handling defects (12). Currently, direct evidence linking specific *POPDC1* frameshift variants to malignant ventricular arrhythmias in a translational model is limited.

To address this gap, we investigated a recessive *POPDC1* frameshift variant (c.448delT) identified in a consanguineous family with early-onset ACM and SCD. We aim to elucidate a novel POPDC1 truncation that perturbs cardiac electrophysiology from cell to whole-organ scale. Using an orthologous *Popdc1* knock-in rat model—selected for its fidelity to human repolarization dynamics—we performed in vivo and ex vivo electrophysiology with biophysical, proteomic, and structural analyses. Our work delineates pathogenic mechanisms linking POPDC1 loss to destabilize the Ankyrin-G (AnkG) scaffold and multi-ion channel remodeling, establishing a potentially lethal re-entry substrate and malignant ventricular arrhythmia.

## Materials and Methods

### Human Subjects and Genetic Analysis

The study protocol adhered to the principles of the Declaration of Helsinki and was approved by the Ethics Committee of Sichuan Provincial People’s Hospital (ID # 2017-114). Written informed consent was obtained from all participants.

We recruited a consanguineous family presenting with early-onset arrhythmia and cardiomyopathy. Genomic DNA was extracted from peripheral blood of the proband, parents, and the affected sister. Whole-exome sequencing (WES) was performed to identify candidate variants, using a capture kit (Agilent SureSelect Human All Exon V6) followed by sequencing on an Illumina platform. Variants were filtered based on minor allele frequency (MAF<0.01 in gnomAD), predicted deleterious effects (loss-of-function), and segregation consistent with autosomal recessive inheritance (13). The candidate *POPDC1* variant was validated and assessed for co-segregation in six extended family members using Sanger sequencing.

### Generation of Popdc1 Knock-in Rats

An orthologous *Popdc1* c.448delT knock-in rat line was generated on the Sprague-Dawley background using CRISPR/Cas9 gene editing (Biocytogen, Beijing, China). A single nucleotide deletion (c.448delT) was introduced into exon 3 of the rat *Popdc1* gene, creating a frameshift (p.Cys150Valfs) analogous to the identified human mutation. This frameshift is predicted to generate a C-terminally truncated BVES protein lacking the functional Popeye domain. Genotypes were identified by PCR, confirmed by Sanger sequencing and Southern blotting (**Figure S1A-D**).

Homozygous mutants (*Popdc1*^fs/fs^) and wild-type littermates (*Popdc1*^+/+^, WT) of both sexes (8-12 weeks; 250-350 g) were used for all experiments, unless otherwise indicated. Animals were housed under controlled environmental conditions (23±1°C; 12/12-h light-dark cycle) with ad libitum access to food and water. All animal procedures were approved by the Animal Care and Use Committee of Chengdu Medical College (ID #2019-186) and conformed to the Guide for the Care and Use of Laboratory Animals (8th Edition, NRC 2011) (14).

### Transthoracic echocardiography

Rats were anesthetized with 4% isoflurane (induction) and maintained with 2% isoflurane for echocardiography (Vevo 3100 LT, FUJIFILM VisualSonics, Canada), with the heart visualized from the parasternal long-axis and short-axis views. The left ventricle ejection fraction (EF) and fractional shortening (FS) were calculated from the measurements of wall thickness and chamber diameters. Left ventricle posterior wall thickness (LVPW), interventricular septal thickness (IVS), stroke volume (SV), and cardiac output (CO) were measured using M-mode. LV mass, ejection fraction (LVEF), and fractional shortening (FS) were calculated using standard formulas (15).

### Electrocardiogram measurements with biotelemetry

Surgery was performed aseptically on rats anesthetized with pentobarbital sodium (40 mg/kg). A catheter-transmitter (Data Sciences International, St. Paul, MN, United States) was inserted into the abdominal wall for ECG monitoring, and the body of the transmitter was implanted in the abdominal cavity. Post-surgery, rats were individually housed and inspected daily for recovery. After two weeks, ECGs of freely moving rats were acquired and ECG traces were analyzed at approximately 10 AM using LabChart v8.2.3 (AD Instruments, Australia). Twelve consecutive complexes were averaged for the reporting of cardiovascular variables. The corrected QT interval (QTc) was calculated using Bazett’s formula (16) unless otherwise indicated. In a subset of animals, rats were subjected to swimming for 15 minutes and ECGs compared before and after exercise.

### In vivo electrophysiology study and arrhythmia induction

Rats were anesthetized with ethyl carbamate (Urethane; KELUN, Chengdu, China) to preserve autonomic function (17). Following tracheal cannulation, animals were mechanically ventilated for electrophysiology study as previously described (18). Briefly, lead II ECG was recorded via subdermal electrodes, with the right femoral vein and artery cannulated for saline infusion and blood pressure monitoring, respectively. A 3F-electrode catheter was advanced via the right jugular vein into the right atrium for programmed electrical stimulation (PES). Atrial/ventricular effective refractory periods (ERP) at basic cycle lengths (BCL) of 150, 120, and/or 100 ms were measured using an S1(×8)-S2 drive train. Ventricular tachycardia (VT) or fibrillation (VF) was defined as >3 consecutive premature ventricular contractions during ERP testing. When non-inducible, double (S3) and triple (S4) extrastimuli were introduced with decremental coupling intervals (starting at 80 ms) until refractoriness. Ventricular arrhythmia inducibility was quantified using a modified arrhythmogenicity index (AI) (19). To assess substrate-dependent repolarization reserve, the non-selective potassium channel blocker cesium chloride (CsCl; 2 mL, 1 mM) was administered intravenously during inducibility testing.

### Hemodynamic assessment

Briefly, rats were anesthetized with ethyl carbamate, mechanically ventilated and surface ECG recorded as described above. A pig-tail catheter was inserted through the right carotid artery into the heart for measuring the left ventricular pressure using LabChart v8.2.3 (ADInstruments, Australia).

### Ex Vivo optical mapping

Rats were anesthetized (pentobarbital sodium, 50 mg/kg i.p.), with hearts were rapidly excised for Langendorff-perfusion with oxygenated Tyrode’s solution (37°C; constant flow: 8 mL/min). Contraction was uncoupled using blebbistatin (10 μmol/L) with hearts stained with voltage-sensitive dye RH237 (1 μg/mL) and excited by 530-nm LEDs (MappingLab Ltd., UK). Fluorescent signals were acquired at 900 frames/s using a CMOS camera system (OMS-PCIE-2002). Data was processed using OMapScope 5.0 to generate activation isochrones and action potential duration (APD) maps (APD20-90). Conduction velocity (CV) and repolarization dispersion were quantified as previously described (20).

### Cellular electrophysiology and calcium imaging

Isolation of ventricular myocytes: Single ventricular myocytes were isolated from rat hearts via enzymatic dissociation as previously described (21).

Whole-Cell patch clamp: Ionic currents were recorded at 37°C using the whole-cell patch-clamp technique with an Axopatch 200B amplifier (Molecular Devices), with pipette resistance at 2-3 MΩ. Peak sodium current (*I*_Na_), L-type calcium current (*I*_Ca,L_), and transient outward potassium current (*I*_to_) were elicited using specific voltage-step protocols as previously described (22), with current identities confirmed using tetrodotoxin (30 µM) and nifedipine (10 µM) in representative cells. Data was analyzed using Clampfit 10.7.

Intracellular Ca² imaging: Cardiomyocytes were loaded with the Ca² indicator Fluo-4 AM (5 μM; Invitrogen) for 30 min at room temperature. Cells were superfused with Tyrode’s solution (1.8 mM Ca²□) at 37°C and field-stimulated at 1 Hz. Spontaneous Ca²□ sparks and waves were recorded using a laser scanning confocal microscope (Leica SP8) in line-scan mode (excitation 488 nm, emission 505 nm). Spark frequency and wave incidence were quantified using ImageJ with the SparkMaster plugin (23).

### Histology and Immunofluorescence

Paraffin-embedded heart sections (5 µm) were stained with Masson’s trichrome and picrosirius red to assess interstitial fibrosis. For immunofluorescence, sections were pre-treated with heat-mediated antigen retrieval (citrate buffer) followed by incubation with primary antibodies against POPDC1 (1:100; Thermo Fisher Scientific; PA5-102304) and AnkG (1:100; Santa Cruz; sc-12719) and respective Alexa Fluor-conjugated secondary antibodies with DAPI counterstaining. *Transmission Electron Microscopy (TEM)*

Myocardial blocks (1 mm³) were fixed in 2.5% glutaraldehyde and 1% osmium tetroxide, dehydrated in graded alcohols, and embedded in Epon 812. Ultrathin sections (70-90 nm) were stained with uranyl acetate and lead citrate. Images were acquired using a HITACHI HT7800 at 80 kV to visualize intercalated disc nanostructures and mitochondrial morphology.

### Molecular and biochemical analysis

Label-Free quantitative proteomics: LV tissue lysates were digested with trypsin and analyzed by LC-MS/MS on a Q Exactive HF-X mass spectrometer (Thermo Fisher Scientific). Raw data were processed using MaxQuant software for protein identification and label-free quantification (LFQ). Differentially expressed proteins were identified based on a fold change>1.5 and P<0.05, and enrichment analyses were conducted on the DEPs using GSEApy (24) with the enriched model.

RT-qPCR and Western blotting: LV tissue lysates were portioned and separately processed for total RNA and protein extracts. Total RNA was reverse-transcribed into cDNA before subjected to quantitative real-time PCR (RT-qPCR) using specific primers listed in **Table S1**, with gene expression levels normalized to β-actin. Protein lysates were separated by SDS-PAGE and transferred to PVDF membranes. Blots were probed with primary antibodies against Cav1.2 (1:1000, MA5-45389, Thermo Fisher Scientific), Kv4.3 (1:500, PA5-93292, Thermo Fisher Scientific), Nav1.5 (1:200, ASC-005, Alomone Labs), TREK-1/KCNK2 (1:1000, PA5-115452, Thermo Fisher Scientific), AnkG (1:1000, 27980-1-AP, Proteintech), BVES (1:1000, as above, Thermo Fisher Scientific), and GAPDH (1:10000, 60004-1-Ig, Proteintech).

Oxidative stress assays: Superoxide dismutase (SOD) activity and malondialdehyde (MDA) content in LV tissue were measured using colorimetric assay kits (Beyotime, China) according to the manufacturer’s instructions.

### Statistical analysis

Data are presented as mean ± SD, unless otherwise stated. Normality was verified using the Shapiro-Wilk test. Comparisons between two groups were performed using unpaired Student’s t-test or Welch’s t-test (for unequal variances). Categorical variables were compared using Fisher’s exact test. For cellular experiments involving multiple cells derived from the same animal, hierarchical data were analyzed using linear mixed-effects models (with animal ID as a random effect) to avoid pseudoreplication (25). All analyses were performed using SPSS 26.0 and GraphPad Prism 9.0. Statistical significance was established at P<0.05.

## Results

### 1. Clinical phenotype and identification of a homozygous POPDC1 frameshift

We identified a consanguineous family with an early-onset, lethal arrhythmogenic cardiomyopathy consistent with autosomal-recessive inheritance (**Figure 1A**). Both parents and one heterozygous sibling were asymptomatic, whereas all three homozygous siblings exhibited a severe, concordant cardiac phenotype.

**Figure 1.**
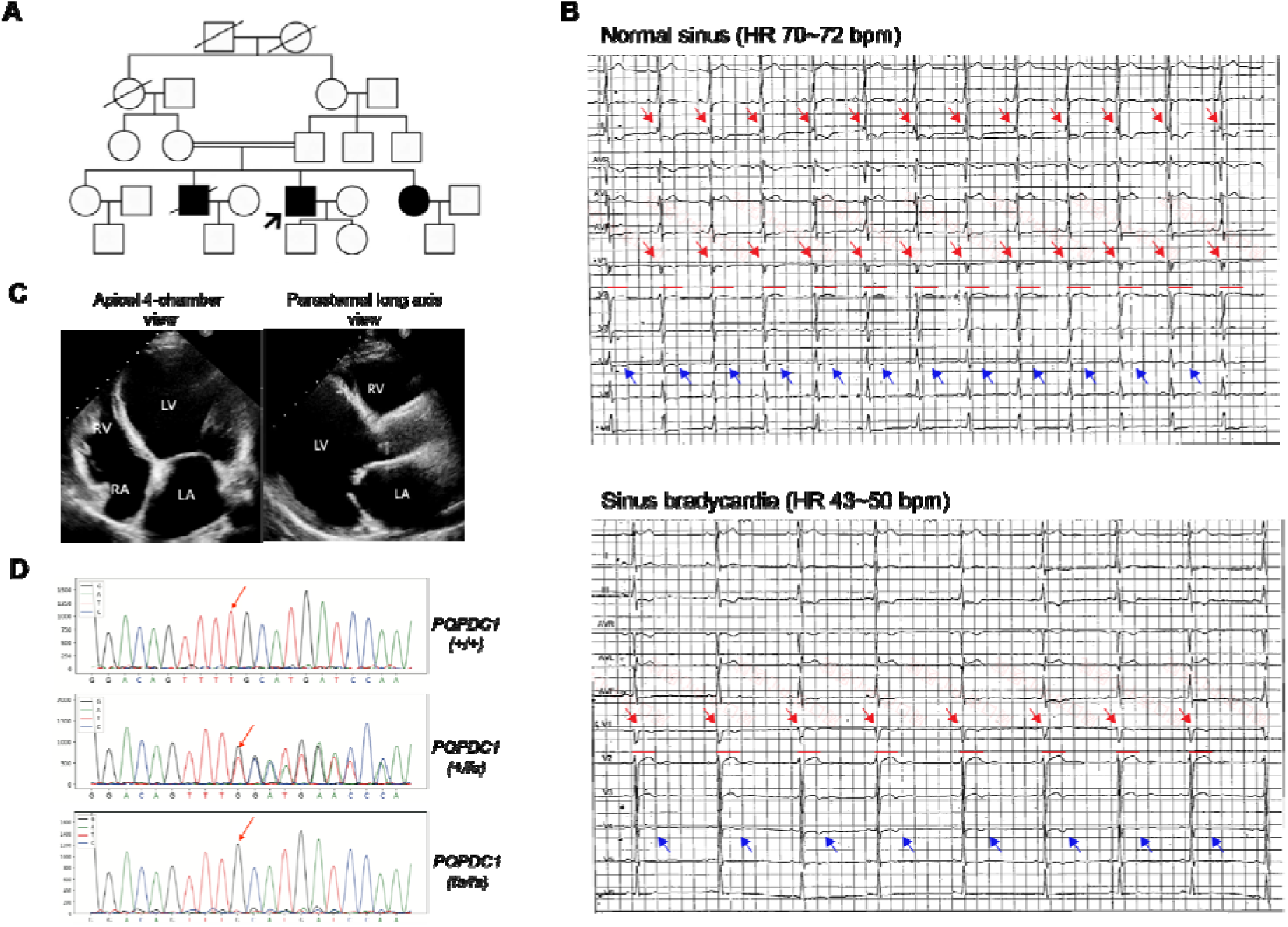
Homozygous POPDC1 truncation causes a lethal arrhythmia-induced cardiomyopathy phenotype. (A) Pedigree of a consanguineous family exhibiting autosomal-recessive inheritance. Solid symbols denote affected individuals presenting with the lethal triad of cardiomyopathy, conduction block, and malignant arrhythmias. Strike-through symbols denote deceased individuals. (B) Representative 12-lead ECG from the proband (age 22) showing normal sinus (top) and sinus bradycardia (bottom) with rate-dependent short QTc interval with evidence of “notched” QRS suggestive of an exhaustion of the repolarization rate-adaptation reserve. Red lines indicate QT interval on lead V2, red arrows indicate notched QRS on leads III and V1 and blow arrows indicate T-wave flattening or inversion on lead V4. (C) Echocardiography demonstrates the late-stage structural phenotype: Dilated cardiomyopathy (DCM)-like characterized by significant LV chamber enlargement and reduced ejection fraction (LVEF). (D) Sanger sequencing validation of the *POPDC1* c.448delT variant. The proband and affected siblings segregate for the homozygous frameshift (fs/fs), whereas parents are heterozygous carriers (+/fs). Red arrow denotes the mutation site. Abbreviations: AV, atrioventricular; ECG, electrocardiogram; LV, left ventricle; LVEF, left ventricular ejection fraction.

The proband’s clinical course displayed an “arrhythmia-first, cardiomyopathy-later” trajectory. At age 18, he presented clinically with exertional dyspnea and binodal dysfunction, including sinus bradycardia (50 bpm) and intermittent first- to second-degree AV block. Dynamic ECG monitoring characterized a pathological maladaptation of the QT-RR relationship. At normal sinus heart rates (70-72 bpm), QTc intervals were preserved within physiological limits (378-416 ms) across four standard correction algorithms (Bazett, Fridericia, Framingham, and Hodges). However, during bradycardic excursions (43-50 bpm), the absolute QT interval exhibited a ‘rate-invariant’ behavior, failing to prolong adequately (persisting at 360–380 ms). This repolarization rigidity precipitated a paradoxical shortening of the QTc to diagnostic short-QT levels (299-363 ms; **Table S2**). This maladaptive repolarization was accompanied by diffuse, rate-independent diffuse T-wave flattening or inversion. Conversely, depolarization abnormalities were rate-dependent. QRS fractionation (notching) emerged during faster heart rates and resolved during bradycardia, indicating exhausted conduction reserve (**Figure 1B**).

Despite dual-chamber pacing, the proband progressed to dilated cardiomyopathy (DCM) -like stage (LVEDD 58 mm, LVEF 43%) at age 22 years (**Figure 1C**). Cardiac resynchronization therapy (CRT-P) failed to halt deterioration, and by age 25 he developed end-stage biventricular failure (LVEDD 81 mm, LVEF 22%; NT-proBNP >18,000 pg/ml) with cardio-hepatic syndrome. Persistent electrical instability (recurrent NSVT and atrial flutter) culminated in SCD three years post-device implantation.

The affected siblings followed a similar course. The older brother developed end-stage heart failure at age 25, underwent transplantation, and died two years later. The affected sister initially exhibited frequent supraventricular ectopy and AV conduction disease, normal sinus with borderline QT interval (360-380 ms) and diffused T-wave flattening/inversion, later progressing to overt LV dilation (LVEDD 58 mm) and systolic dysfunction (LVEF 44%). None of the homozygous siblings showed clinical myopathy, although serum creatine kinase (CK) was mild-to-moderately elevated (500–1000 IU/L) in the proband and his affected sister. This combination of lethal arrhythmic cardiomyopathy with subclinical skeletal involvement contrasts with classical LGMDR25 (4, 6) and may support a primary diagnosis of ACM rather than secondary DCM.

### 2. Genetic analysis identify a pathogenic POPDC1 truncation

Whole-exome sequencing was performed on the proband and nuclear family to define the genetic etiology. Variants in established arrhythmia- and cardiomyopathy-associated genes, including *KCNH2*, *KCNQ1*, *SCN5A*, *LMNA*, and desmosomal genes, were first excluded. A stepwise filtering pipeline prioritizing rare, protein-altering variants under an autosomal-recessive model yielded five candidates. Segregation analysis by Sanger sequencing in six extended family members demonstrated that only variants in *POPDC1* and *SERINC1* co-segregated with the phenotype; *POPDC1* was prioritized for its cardiac expression and function. The homozygous *POPDC1* c.448delT variant was absent from public databases and this single-nucleotide deletion introduces a frameshift (p.Cys150Valfs) and a premature stop codon at position 151, producing a truncated BVES protein lacking the conserved Popeye domain essential for cAMP binding (**Figure 1D**). The predicted loss-of-function mechanism (PVS1), combined with the absence from population databases (PM2) and strong segregation evidence (PP1), supports classification of the variant as Pathogenic under ACMG/AMP criteria (26).

Screening of sporadic early onset atrial fibrillation (AF) (n=222) and DCM (n=344) cohorts identified rare heterozygous POPDC1 variants (<1%). An AF carrier (p.Glu337) exhibited J-wave elevation and bradyarrhythmia requiring pacemaker implantation, while a DCM carrier (p.Lys214Asn) experienced sudden cardiac death at age 50 (**Table S3)**. These heterozygous presentations were markedly milder and later in onset than the catastrophic early lethality (<30 years) observed in the above homozygous pedigree.

### 3. Longitudinal characterization reveals stage-dependent structural remodeling in Popdc1^fs/fs^ rats

To model the human disease, we generated an orthologous *Popdc1* c.448delT knock-in rat (**Figure S1A-D**). Homozygous (*Popdc1*^fs/fs^) and heterozygous (*Popdc1*^fs/+^) animals were viable, fertile, and exhibited normal growth compared with WT littermates (*Popdc1*^+/+^). Consistent with the recessive inheritance in the clinical family, heterozygous rats displayed no ECG or echocardiographic abnormalities and were not studied further.

Longitudinal assessment showed clear stage-dependent remodeling. Echocardiography was unremarkable at approximately 2-months, but by 3–4 months *Popdc1*^fs/fs^ rats developed pronounced LV wall thickening (hypertrophy) and hypercontractility (**Figure S2A-C**). Cardiac hemodynamics in 3-4 month-old adult rats demonstrated comparable LV end-systolic pressure (LVESP) and heart rates between WT and *Popdc1*^fs/fs^ rats, indicating preserved systolic function despite an emerging HCM-like phenotype (**Figure S3A**), and histology revealed marked interstitial fibrosis (**Figure S3B**).

### 4. Increased susceptibility to AV-block in Popdc1^fs/fs^ rats

To determine whether electrical instability precedes structural remodeling, we evaluated *Popdc1*^fs/fs^ rats at the pre-structural stage (2-3 months). Baseline telemetry confirmed P-wave prolongation and shortened QT, QTc, and J-Tpeak intervals relative to WT (**Figure 2A-F**). Upon swimming stress, *Popdc1*^fs/fs^ rats exhibited chronotropic incompetence, failing to generate the tachycardic response seen in WT controls, instead revealing exhaustion of the conduction reserve: a blunted heart-rate response and progressive, rate-dependent PR prolongation (**Figure 2G-J**) and ECG demonstrated repolarization abnormalities, including shortened QT/QTc interval and diffuse T-wave flattening or inversion (**Figure 2G-J; Figure S4A-C**).

**Figure 2.**
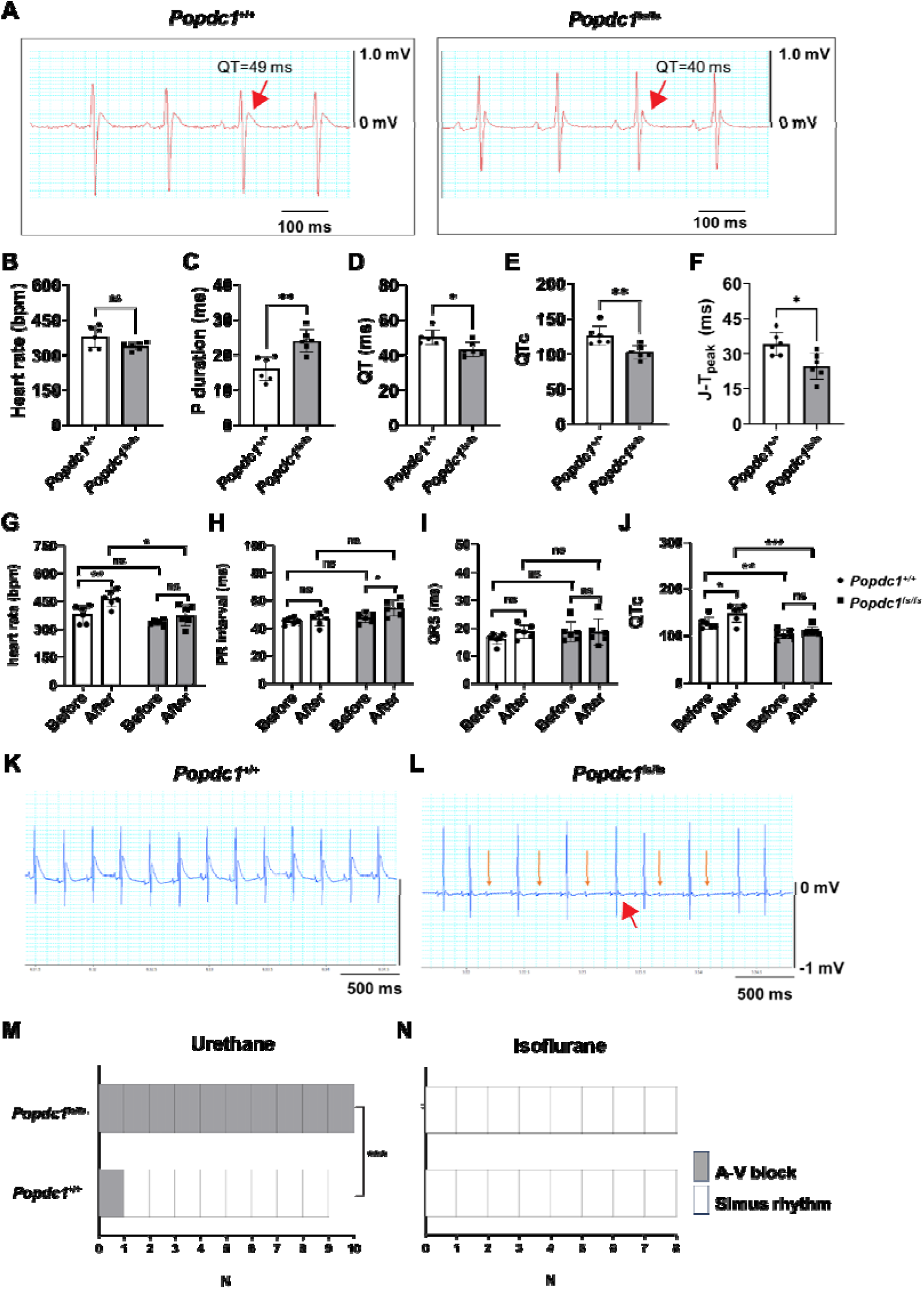
Pharmacological unmasking reveals a critical conduction-repolarization mismatch in *Popdc1*^fs/fs^ rats. (A) Representative telemetry ECG traces from conscious *Popdc1*^+/+^ (WT) and *Popdc1*^fs/fs^ rats. Red arrows show the calculated QT intervals demonstrating a distinctive shortened QT in *Popdc1*^fs/fs^ rats (40 ms vs 49 ms in WT). (B-F) Calculated heart rate (B), P duration (C), QT interval (D), corrected QT (E), and J-T peak interval (F) demonstrating abnormal conduction in *Popdc1*^fs/fs^ rats (n=6/group). Summary statistics reveal a specific “slow-conduction, short-refractory” profile: prolonged P-wave duration (conduction slowing) coupled with shortened QT, QTc, and J-Tpeak intervals under preserved basal heart rate. (G-J) Differences in heart rate (G), PR interval (H), QRS (I), and corrected QT (J) in WT and *Popdc1*^fs/fs^ rats before and after swimming (n=6/group). (K-N) Representative ECG trace during urethane anesthesia (a mild Na^+^ channel suppressor) in controls (K) and *Popdc1*^fs/fs^ rats (L). Red and orange arrows indicate a T-wave depression and high-grade AV block in the representative *Popdc1*^fs/fs^ rat, absent in controls. (M-N) Proportion of rats with AV-block or sinus rhythm under urethane (M) or isoflurane (N) anesthesia (n=8-10/group). Contrastingly, isoflurane anesthesia preserves 1:1 conduction. This “state-dependence” confirms a latent frailty in sodium channel function. Data are presented as mean ± SD (B-J) and proportions (M-N). Statistical analyses were conducted with unpaired Student’s t-test (B-F), paired t-test (G-J), and Fisher’s exact test (M-N). *P<0.05, **P<0.01, ***P<0.001, ns = not significant.

Under urethane anesthesia, which mildly suppresses sodium currents (27), all *Popdc1*^fs/fs^ rats exhibited susceptibility to second-degree AV block and junctional escape rhythms compared to 1 of 9 controls (**Figure 2K-M**). Conversely, isoflurane which modulates electrophysiology via enhancement of outward potassium currents (28), with relatively limited direct inhibition of sodium channels at clinically relevant concentrations, did not induce AV block in either genotype (**Figure 2N**). Like the clinical proband, *Popdc1*^fs/fs^ rats frequently exhibited transient T-wave flattening or inversion (**Figure 2L**), providing further support of a conserved, stress-exacerbated repolarization defect. This suggests that the conduction abnormality was unmasked specifically under sodium channel-challenging conditions.

### 5. VT susceptibility in Popdc1^fs/fs^ rats

Consistent with the short-QT phenotype on surface ECG, *Popdc1*^fs/fs^ rats exhibited tendencies for atrial (AT/AF) and ventricular arrhythmias (VT) with PES (**Figure 3A-D**), coinciding with a global abbreviation of ERP. Both atrial ERP (AERP) and ventricular ERP (VERP) were significantly reduced compared to WT littermates across all pacing cycle lengths (150, 120, and/or 100 ms). (**Figure 3E-F**).

**Figure 3.**
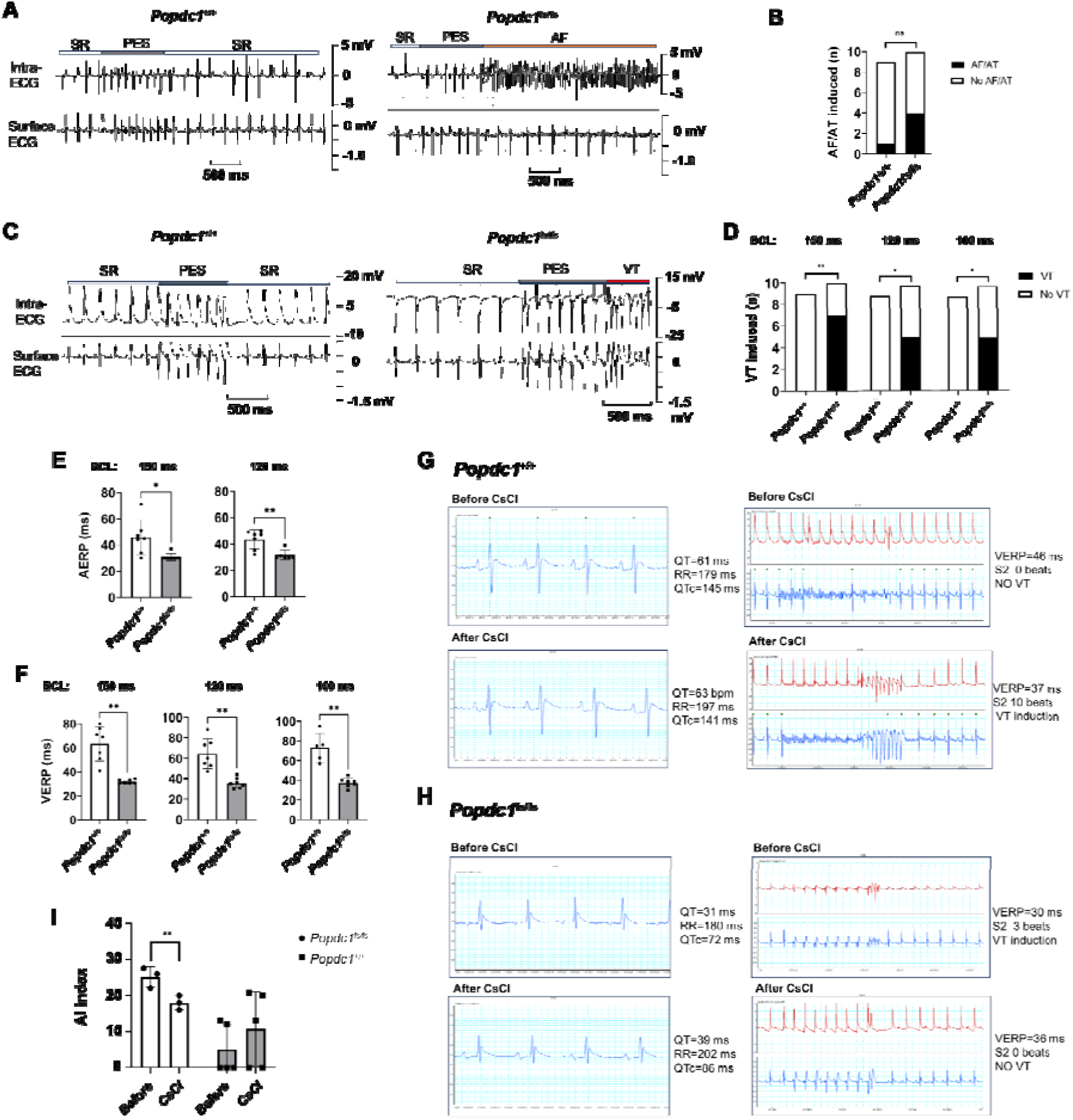
High arrhythmia inducibility and global refractory period abbreviation confirm a re-entrant substrate in *Popdc1*^fs/fs^ rats. Representative intracardiac electrograms during programmed electrical stimulation (A) demonstrating increased atrial tachycardia/atrial fibrillation (AF/AT) induction in *Popdc1^fs/fs^* hearts (B). Representative intracardiac electrograms during programmed electrical stimulation (C) demonstrating high vulnerability to ventricular tachycardia (VT) induction in *Popdc1^fs/fs^* hearts (D). (E-F) Global refractoriness abbreviation evident by atrial (AERP; E) and ventricular effective refractory periods (VERP; F) as measured under basic cycle lengths (BCL) of 150, 120, and/or 100 ms intervals, confirming an intrinsic hyper-repolarization defect.. (G-H) Representative traces demonstrate that cesium chloride (CsCl) administration significantly prolongs the QT interval and suppresses arrhythmias in WT (B) compared to *Popdc1*^fs/fs^ rats (C), suggesting a “repolarization gain-of-function” affecting potassium currents in the mutants. This “rescue effect” mechanistically identifies excessive potassium currents as the driver of instability. (D-F) Proportion of rats per group with demonstrable VT measured at basic cycle lengths of 150, 120, and 100 ms. Sample sizes as indicated for A-F (n=9-10/group), E-F (n=5-7/group), and I (n=3-5/group). Data are presented as proportions (B, D) and mean ± SD (E, F, and I). Statistical analyses were conducted with Fisher’s exact test (B, D), unpaired Student’s t-test (E-F), paired t-test (I). *P<0.05, **P<0.01 vs. WT.

To further probe the electrophysiological substrate, CsCl (inhibiting background potassium currents) was administered to modulate repolarizing currents. In control rats, CsCl tended to increase AI consistent with its pro-arrhythmic effects, whereas AI was lowered in *Popdc1*^fs/fs^ rats (**Figure 3G-I**).

### 6. Optical mapping identifies a coupled slowed conduction-abbreviated repolarization substrate

To decipher the tissue basis for the observation of “conduction-repolarization mismatch” (P-wave prolongation with short-QT/QTc; **Figure 2C-F**), we performed high-resolution optical mapping of the ventricular activation (**Figure 4A**). Isochronal mapping demonstrated markedly delayed propagation across *Popdc1*^fs/fs^ hearts, reflecting a significant reduction in CV (**Figure 4B, C**). Although time-to-peak activation remained preserved (**Figure 4D-F**), action potential amplitude (APA) was significantly reduced in *Popdc1*^fs/fs^ hearts (**Figure 4G**). This tissue-level loss of APA aligns with the in vivo urethane-unmasked phenotype of AV-block (**Figure 2K-M**), confirming that conduction slowing arises from reduced cellular excitability, likely diminished *I*_Na_, rather than impaired intercellular coupling.

**Figure 4.**
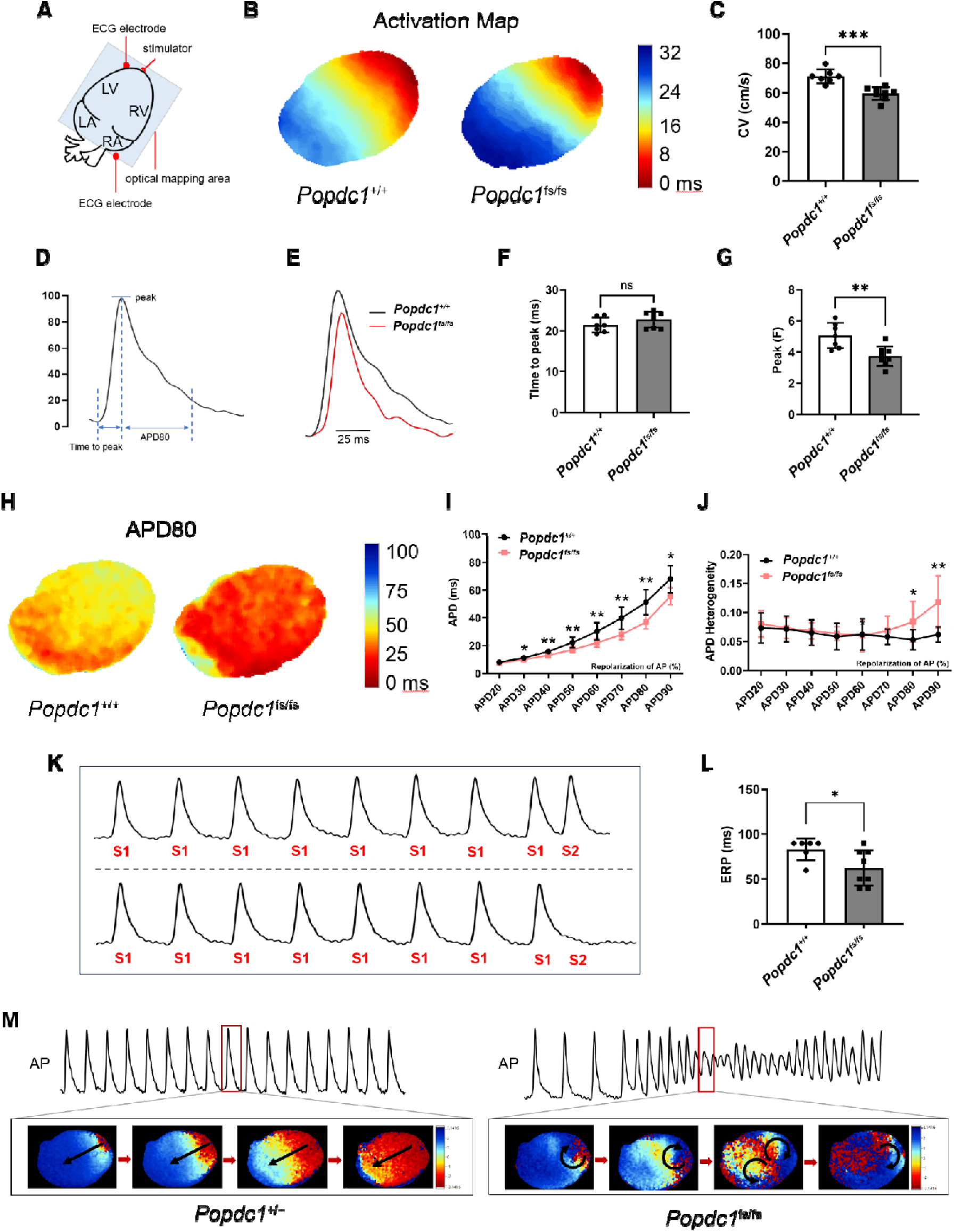
Optical mapping deciphers the biophysical substrate in *Popdc1*^fs/fs^ rats: slowed conduction coupled with accelerated repolarization. (A) Schematic of the ex vivo Langendorff perfusion and optical mapping setup. (B–C) Representative isochronal maps (B) reveal delayed activation times in *Popdc1*^fs/fs^ hearts reflected by slowed average conduction velocity (CV; C). (D–G) Example action potential (D) depicting AP indices of time-to-peak (depolarization time), peak amplitude, and APD80 (repolarization time) measurements as indicators of cellular excitability. (E) Representative optical action potential (OAP) traces in *Popdc1*^fs/fs^ and *Popdc1*^+/+^hearts. While time-to-peak is preserved (F), the AP amplitude (Peak) is significantly reduced (G), suggestive of tissue-level evidence of sodium channel dysfunction. (H-J) Representative APD80 maps (H) demonstrating wide-spread action potential duration (APD) abbreviation across all phases of repolarization (APD20–90; I) and inhomogeneity in late-phase APD repolarization in the mutants (J), potentially favoring unidirectional block. (K–L) Ex vivo effective refractory period (ERP) quantification by S1-S2 PES protocol (K) confirms shortened ERP in mutants compared to controls (L). (M) Representative phase singularity maps visualizing stable re-entrant rotors (spiral waves) that sustain polymorphic VT in the mutants compared to planar conduction in the WT controls. Sample sizes as indicated were n=7-8/group (B-J) and n=6-8/group (L). Data expressed as mean ± SD. Statistical analyses by unpaired Student’s t-test (B, F, G), two-way ANOVA with (I,J). *P<0.05, **P<0.01, ***P<0.001 respectively.

Consistent with the short-QT phenotype, optical action potentials (OAPs) showed global repolarization abbreviation in *Popdc1*^fs/fs^ hearts. APD80 (**Figure 4H-J**) and ERP were significantly shortened (**Figure 4K, L**), and this reduction spanned all repolarization phases (APD20–APD90), indicating a broad acceleration of repolarizing currents that mirrors the cycle-length-independent ERP shortening observed during PES (**Figure 3E, F**). Spatial repolarization heterogeneity was increased at later phases (80-90%) of the APD (**Figure 4J**), creating a substrate permissive to unidirectional block and arrhythmia.

PES utilizing an S1-S2 protocol readily induced sustained polymorphic ventricular tachycardia (PVT) with a torsade-like morphology in one isolated *Popdc1*^fs/fs^ hearts. Optical mapping revealed stable re-entrant rotors sustaining arrhythmia (**Figure 4M**). Because arrhythmias were only triggered by premature S2 beats, these findings implicate calcium-dependent triggered activity as the initiating event that ignites a highly vulnerable, short-wavelength re-entrant substrate.

### 7. Cellular ionic mechanisms underlying the pro-arrhythmic substrate

To dissect the ionic basis of the “conduction-repolarization mismatch” observed in vivo and ex vivo, we further evaluated major currents in isolated ventricular myocytes. Guided by the optical mapping signature—blunted APA and globally abbreviated and heterogeneous duration (**Figure 4D-J**)—we focused on the principal currents governing depolarization (*I*_Na_), early repolarization (*I*_to_), and calcium-dependent triggering (*I*_Ca,L_).

Patch-clamp recordings revealed a striking loss-of-function in the fast sodium current (*I*_Na_). Peak *I*_Na_ density in *Popdc1*^fs/fs^ myocytes was markedly reduced by 65% at −30 mV compared to controls (P<0.01, **Figure 5A-C**). This deficit potentially explains the slowed CV and reduced APA observed optically (**Figure 4C, G**), as well as the urethane-induced conduction failure (**Figure 2M**).

**Figure 5.**
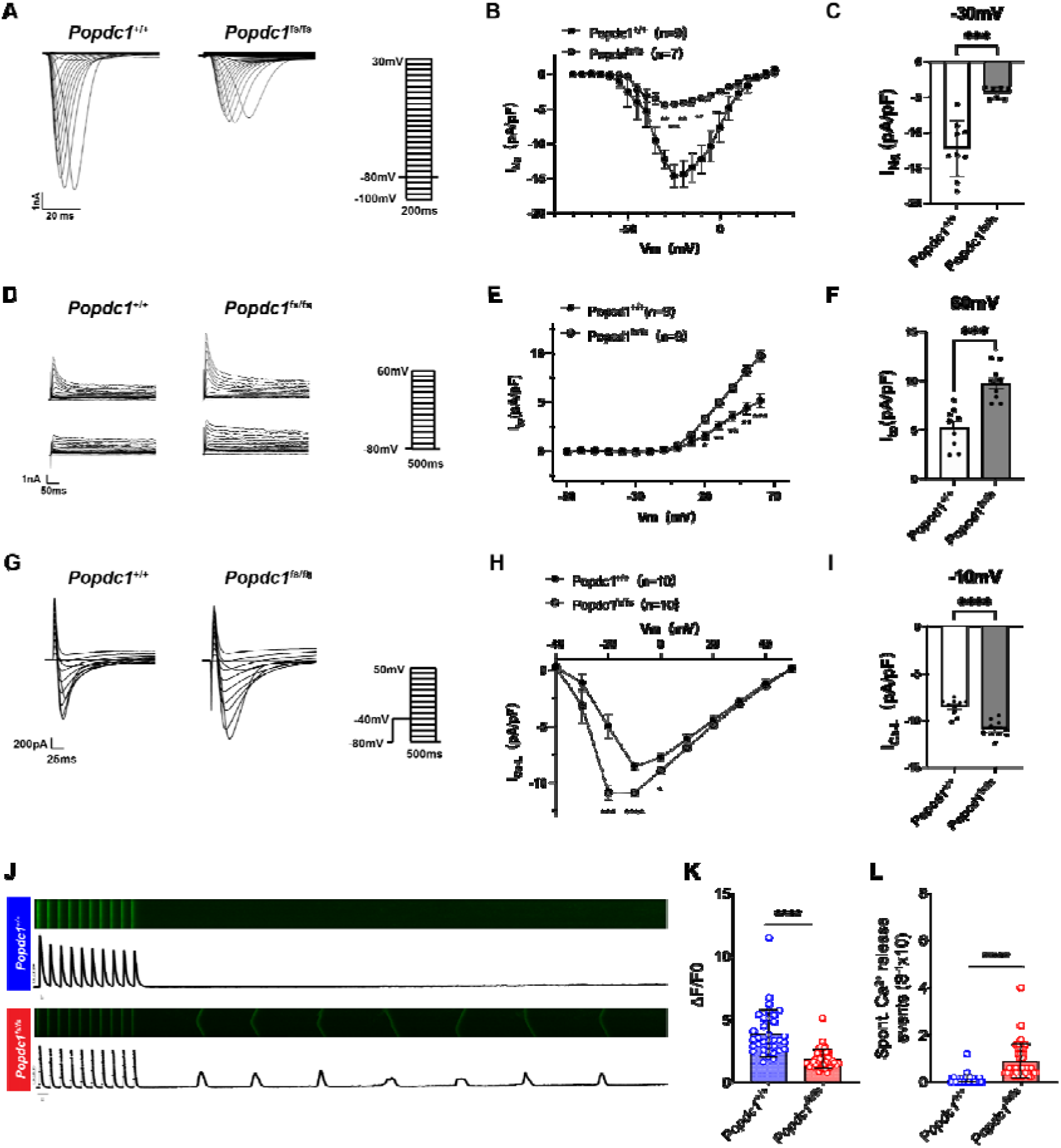
Altered ion channel remodeling and calcium instability in *Popdc1*^fs/fs^ rats drives the cellular arrhythmogenic mechanisms. (A–I) Representative recordings of sodium (*I*_Na_, A), transient outward potassium (*I*_to_, D) and L-type calcium (*I*_Ca,L_, G) currents in *Popdc1*^+/+^ (WT controls) and *Popdc1*^fs/fs^ cardiomyocytes. Current-voltage relationships across step-wise voltage-clamp *I*_Na_ (B), increased *I*_to_ (E), and *I*_Ca,L_ (H) demonstrate the divergence of peak current amplitudes, showing reduced peak current amplitudes of *I*_Na_ (C), increased *I*_to_ (F), and *I*_Ca,L_ (I) in *Popdc1*^fs/fs^ cardiomyocytes (n=9-10/group). (I-J) Confocal line-scans (I) and quantification (J) reveal a significantly higher incidence of spontaneous Ca^2+^ waves in *Popdc1*^fs/fs^ myocytes, consistent with sarcoplasmic reticulum instability (n=3-4/group (3-5 cardiomyocytes per rat)). Data are mean ± SEM. Statistical analysis by two-way ANOVA with Tukey post-hoc test (B, E, H), unpaired Student’s t-test (C, F, I, K, L). *P<0.05, **P<0.01, ***P<0.001, ****P<0.0001 respectively.

Conversely, the transient outward potassium current (*I*_to_) exhibited a robust gain-of-function. Peak *I*_to_ density was significantly increased in *Popdc1*^fs/fs^ myocytes, particularly at positive potentials (P<0.0001 at +60 mV) (**Figure 5D-F**). This intrinsic acceleration of repolarization aligns with the CsCl “rescue” effect seen in telemetry studies (**Figure 3I**) and accounts for the cycle-length-independent ERP shortening during PES (**Figure 3E, F**).

To identify the trigger for arrhythmia initiation, we quantified L-type calcium current. *I*_Ca,L_ density was markedly elevated in *Popdc1*^fs/fs^ myocytes (P<0.0001 at −10 mV) (**Figure 5G-I**). Confocal calcium imaging confirmed that this enhanced influx destabilizes the sarcoplasmic reticulum (SR) calcium handling, producing a significantly higher frequency of spontaneous Ca^2+^ waves (**Figure 5J-L**). These stochastic release events provide the delayed afterdepolarizations triggers required to ignite arrhythmias on the vulnerable *I*_Na_/*I*_to_-electrophysiological substrate.

Together, the combined *I*_Na_ loss-of-function, *I*_to_ and *I*_Ca,L_ gain-of-function establish a coherent ionic mechanism linking reduced excitability, accelerated repolarization and calcium-dependent triggered activity in *Popdc1*^fs/fs^ myocytes.

### 8. Proteomics reveals post-translational scaffold collapse and bipartite ion channel remodeling

Unbiased label-free quantitative proteomics of *Popdc1*^fs/fs^ ventricular myocardium uncovered a defining molecular signature. Volcano plot analysis identified marked downregulation of BVES (POPDC1) and the membrane cytoskeletal adaptor AnkG as the dominant defects (**Figure 6A**). KEGG enrichment of differentially expressed proteins highlighted “Arrhythmogenic Right Ventricular Cardiomyopathy (ARVC)”, “Dilated Cardiomyopathy (DCM)”, and “Cardiac muscle contraction” pathways (**Figure 6B**), indicating that POPDC1 truncation initiates broad cardiac structural remodeling.

**Figure 6.**
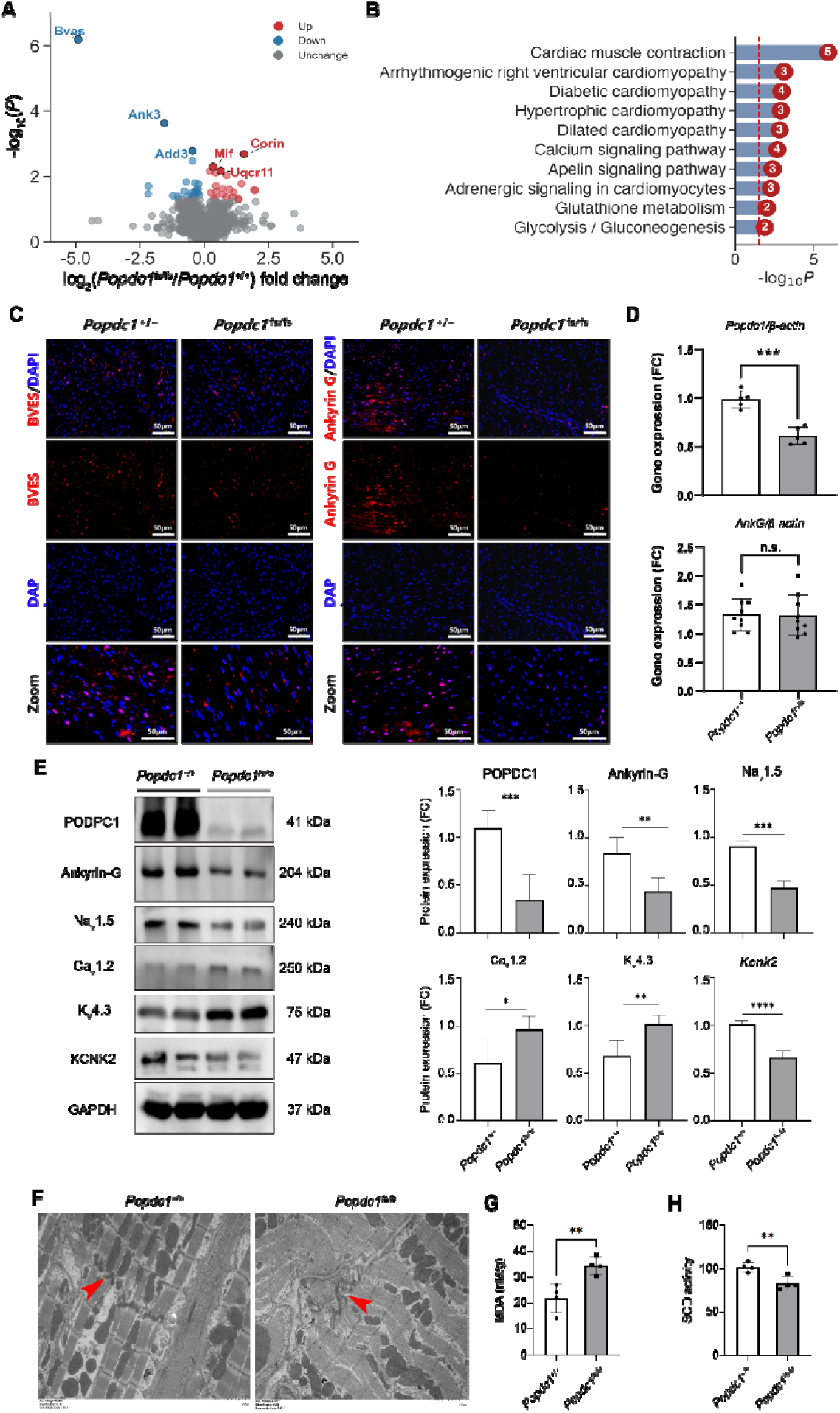
Proteomics and RT-qPCR indicate potential of post-translational scaffold collapse, channel mis-trafficking, and oxidative stress in *Popdc1*^fs/fs^ rats. (A) Volcano plot depiction of the differential proteome from ventricular tissue lysates of *Popdc1*^+/+^ and *Popdc1*^fs/fs^ rats. Prioritization of the top targets highlighted profound downregulation of BVES (POPDC1) and the membrane scaffold Ankyrin-G (Ank3) (n=5-7). (B) KEGG pathway enrichment of the proteome identified “Cardiac muscle contraction” and “Arrhythmogenic right ventricular cardiomyopathy” and various cardiomyopathies as the top disrupted pathways. (C) Immunofluorescence staining of the myocardium demonstrates the loss of both BVES and Ankyrin-G (both in red) in *Popdc1*^fs/fs^ rats. (D) RT-qPCR gene expression analyses in cardiac tissue demonstrated reduced *Popdc1* but normalized *AnkG* mRNA (n=5-7). (E) Western blots confirm the reduction of cardiac POPDC1-Ankyrin-G-Nav1.5 axis and TREK-1 (KCNK2), concurrent with an upregulation of Kv4.3 and Cav1.2 in *Popdc1*^fs/fs^ rats (n=6). (F–G) *Popdc1*^fs/fs^ hearts exhibit markers of oxidative stress: elevated malondialdehyde (MDA; G) and reduced superoxide dismutase (SOD; H) activity (n=4). (F) Transmission electron microscopy (TEM) displays widened intercalated discs and myofibrillar disarray (red arrowhead) in *Popdc1*^fs/fs^ hearts (n=3). Data are mean ± SD. Statistical analyses conducted were unpaired Student’s t-test. *P<0.05, **P<0.01, ***P<0.001, ****P<0.0001 respectively.

To define the mechanism underlying this interactome loss, we integrated spatial and transcriptional analyses. Immunofluorescence established the structural basis, confirming that reduced BVES and AnkG expression in *Popdc1*^fs/fs^ hearts (**Figure 6C**). However, transcriptional profiling by RT-qPCR revealed a key discordance, decreased *Popdc1* mRNA was significantly reduced (P<0.001) but unchanged *Ank3* (AnkG) mRNA (**Figure 6D**) despite profound protein depletion confirmation on Western Blotting (**Figure 6E**). These findings suggest that loss of the POPDC1 anchor potentially drives rapid post-translational degradation of “unmoored” AnkG. Correspondingly, TEM demonstrated widened and disorganized intercalated discs in mutant myocardium (**Figure 6F**).

Consistent with the loss of the AnkG scaffold (29) and POPDC1 chaperone function, Nav1.5 and TREK-1 (KCNK2) were significantly downregulated (**Figure 6E**) (2), confirming that the *I*_Na_ deficit arises from proteostatic instability rather than transcriptional repression. TREK-1 loss likely further promotes steady-state sodium channel inactivation by depolarizing the resting membrane potential (2,4). Conversely, Kv4.3 and Cav1.2 were upregulated (**Figure 6E**), mirroring the short-QT phenotype in *Popdc1*-null zebrafish (12) and implicating Kv4.3 induction as a conserved driver of accelerated repolarization.

This combined ion-channel remodeling converged with a metabolic disturbance to trigger intracellular calcium instability. *Popdc1*^fs/fs^ hearts exhibited a pro-oxidative state evidenced by elevated MDA and reduced SOD activity (**Figure 6G, H**). Oxidative stress sensitizes RyR2 channels [30, 31] and, together with Cav1.2-mediated calcium overload, destabilizes the sarcoplasmic reticulum. Consequently, the heightened frequency of spontaneous Ca^2+^ waves observed in mutant myocytes (**Figure 5J-L**) generates delayed afterdepolarizations, serving as the spark to initiate arrhythmias on the vulnerable, short-wavelength substrate.

## Discussion

### Recessive short-QT ACM as a distinct clinical entity

This study identifies homozygous *POPDC1* truncation as the genetic driver of a previous unrecognized disorder we term rSQT-ACM. Distinct from the classical limb-girdle muscular dystrophy (LGMDR25) phenotype associated with biallelic *POPDC1* variants (4,5), this condition is defined by a characteristic diagnostic triad: (1) early-onset bradyarrhythmia (sinus bradycardia or AV block); (2) a paradoxical short-QT interval with maladaptive rate response; and (3) progressive arrhythmogenic cardiomyopathy accompanied by malignant ventricular arrhythmias. Unlike classic cardiomyopathies where electrical remodeling is secondary to structural failure, rSQT-ACM represents a distinct cardiomyopathy from POPDC1 loss–associated AnkG destabilization and resultant multi-ion channel remodeling.

A defining clinical hallmark is the rigidity repolarization: although QT intervals appear borderline short, they fail to prolong during bradycardia, indicating a constitutive repolarizing drive likely mediated by *I*_to_ upregulation. Pedigree and longitudinal follow-up consistently demonstrate an electrical-to-substrate progression, with conduction disease and repolarization abnormalities preceding ventricular dilation. This sequence establishes electrical instability as the primary etiology, aligning this entity with the emerging framework of arrhythmogenic left ventricular cardiomyopathy (ALVC) (32, 33). Consequently, electrical destabilization acts as the primary driver, challenging the traditional dichotomy between channelopathies and cardiomyopathies.

The electrophysiological signature represents a mechanistic overlap between two malignant channelopathies: *I*_Na_ loss reminiscent of Brugada Syndrome (34), and repolarization gain characteristic of short-QT Syndrome (SQTS) (35, 36). These findings expand the phenotypic spectrum of the POPDC1-related disease beyond muscular dystrophy and supports prioritizing *POPDC1* in the genetic evaluation of young patients with unexplained cardiomyopathy and short-QT intervals—traditionally attributed to primary channelopathies in structurally normal hearts (35).

### Scaffold destabilization and the erosion of conduction reserve

Although POPDC proteins are broadly implicated in membrane trafficking and cAMP signaling (37), our findings reveal a specific structural requirement for POPDC1 in maintaining the intercalated disc (ID) nanodomain. The central defect is a post-translational “scaffold collapse” of the critical regulator of cytoskeletal organization and cardiac integrity, AnkG (38, 39). Despite preserved *Ank3* transcript levels, AnkG protein is profoundly depleted, demonstrating that POPDC1 acts as an essential stabilizing anchor. Loss of this anchor triggers rapid degradation of “unmoored” AnkG, disrupting the master organizer responsible for clustering Nav1.5 at the ID (29).

This collapse produces a hierarchical conduction defect, (1) structural loss of Nav1.5 due to AnkG depletion (38) and (2) functional inactivation of the remaining Nav1.5 pool due to TREK-1/KCNK2 loss and resting membrane potential depolarization (2,4). Together, these mechanisms erode the conduction reserve, explaining the rate-dependent QRS fractionation in the proband and dramatic conduction collapse during physiological stress in the rat model. Mild sodium channel suppression with urethane unmasked high-grade block, confirming that the scaffold-depleted substrate lacks the safety margin required for frequency adaptation and is vulnerable to metabolic-electrical mismatch.

### Opposing remodeling and wavelength collapse

Physiologically, the QT interval prolongs as heart rate slows to maintain a stable QTc. In contrast, our proband exhibited pathological repolarization during bradycardia: the QT interval failed to adapt, resulting in anomalous QTc shortening. This clinical anomaly was validated in the *Popdc1*^fs/fs^ rat, where in vivo ECGs and ex vivo optical mapping confirmed a shortened refractory substrate. Unlike the classic slow conduction/long repolarization remodeling seen in terminal heart failure (40), our patch-clamp analysis revealed a distinct pattern of “opposing remodeling”: severe *I*_Na_ reduction coupled with paradoxical upregulation of Kv4.3-mediated *I*_to_. The latter likely reflects a maladaptive response to oxidative stress, consistent with the susceptibility of potassium channels to the redox signaling (41,42).

This remodeling produces a biophysical paradox. *I*_to_-mediated acceleration of the Phase 1–3 repolarization shortens the ERP, while *I*_Na_ loss slows CV. According to the wavelength (WL) equation (WL = CV x ERP) (43), simultaneous reductions in CV and refractoriness (ERP) causes a collapse of excitation wavelength, enabling re-entry within a small anatomical substrate. This explains the high arrhythmic burden despite the absence of marked ventricular dilation.

The substrate is then triggered by calcium dysregulation and metabolic stress. Increased *I*_Ca,L_ density, spontaneous Ca^2+^ sparks/waves, and a pro-oxidant milieu (elevated MDA, reduced SOD) indicate ROS-mediated RyR2 sensitization (30, 31), providing the delayed afterdepolarizations that ignites re-entry on this short-wavelength substrate. Furthermore, this oxidative insult also converges with AnkG disruption to drive the progressive structural remodeling and cardiomyopathy (31, 38).

### Study limitations and Translational Perspectives

Our study has several limitations. First, although the proband presented with an end-stage DCM-like phenotype, the *Popdc1*^fs/fs^ rats exhibited cardiac hypertrophy and fibrosis despite preserved function. This divergence likely reflects different progression points along the same disease continuum. Specifically, the model captures the critical arrhythmogenic window of ACM—the early phase where electrical instability precedes hemodynamic decline—thereby providing a tractable platform to dissect primary arrhythmic mechanisms independent of the secondary remodeling that accompanies terminal pump failure. Second, species-specific differences in repolarization must be acknowledged. Rodent ventricular repolarization is dominated by the *I*_to_ current, whereas human myocytes rely more on the delayed rectifier currents (*I*_kr_/ *I*_ks_) (44). Despite these ionic distinctions, the overarching principle of “scaffold-mediated repolarization gain”—in which loss of upstream structural regulation drives maladaptive potassium conductance—is likely conserved across species. Future studies utilizing patient-specific iPSC-CMs will be essential to determine whether *I*_kr_/*I*_ks_ remodeling parallels the *I*_to_ augmentation observed here. Finally, the precise molecular interface underlying the functional interdependence within the POPDC1-Ankyrin-G nanodomain remains unresolved. Distinguishing direct physical or intermediary interaction will be a key objective for future structural and biochemical studies.

### Clinical implications and therapeutic framework

Recognizing this disorder as rSQT-ACM has immediate diagnostic and therapeutic implications. The combination of binodal disease (sinus bradycardia or AV block), atypical short-QT intervals, and progressive cardiomyopathy should prompt early POPDC1 sequencing to avoid misclassification as idiopathic DCM.

Mechanistically, the profound depolarizing reserve deficit renders sodium channel blockers contraindicated. Even mild *I*_Na_ suppression in the rat model precipitated high-grade AV block, underscoring the risk of exhausting the already compromised conduction reserve and worsening re-entry by further slowing CV.

Because arrhythmic instability precedes overt pump failure, early ICD implantation may offer superior protection compared with standard bradycardia pacing. Ultimately, defining this pathology as a recessive scaffoldopathy shifts the therapeutic paradigm toward mechanism-based gene replacement strategies aimed at restoring the intercalated disc nanodomain rather than treating downstream electrical or structural sequelae.

## Conclusions

In conclusion, we characterize rSQT-ACM as a lethal entity caused by biallelic *POPDC1* truncation. The resultant loss of Ankyrin-G scaffolding drives a unique “opposing remodeling” phenotype (severe *I*_Na_ reduction coupled with paradoxical upregulation of Kv4.3-mediated *I*_to_), manifesting as bradycardia, paradoxical short-QT intervals, and ventricular tachycardia. These insights dictate immediate clinical changes: the avoidance of sodium channel blockers and the prioritization of early ICD implantation. While validation in larger cohorts is warranted, this distinct triad offers a vital roadmap for identifying and protecting at-risk patients.

## Acknowledgements

**Author Contributions**

X.L. conceptualized the study, analyzed the clinical pedigree data, and critically revised the manuscript. R.L. and X.C. drafted the manuscript. L.Y., X.W., and W.H. contributed to experimental design and data interpretation. R.L. supervised the generation of the point-mutation rat model and characterized the cardiomyopathy phenotype. X.C. and Y.W. performed in vivo electrophysiological studies and formal data analysis. H.L. and Y.H. executed cellular patch-clamp recordings. Q.S. acquired echocardiographic data. S.L. performed optical mapping experiments. H.D. conducted protein and molecular assays. W.L. contributed to figure presentation, revised the manuscript and provided linguistic editing.

## Funding

Financial support was obtained from the National Natural Science Foundation of China (81770379, 81670290); Zhong Nanshan Medical Foundation of Guangdong Province (ZNSA-2020017). Natural Science Foundation of Sichuan Province (2022NSFSC0538). W.L. is supported by the SingHealth Duke-NUS Academic Medicine-Designated Philanthropic Funds (2025/EX/00391).

## Conflict of interest

None declared.

## Data availability

The data sets analyzed in this study are available from the corresponding author upon reasonable request.

## Notes

### Competing Interest Statement

The authors have declared no competing interest.

