## Supplemental for "Recessive *POPDC1* Truncation Causes Lethal Short-QT Pattern Arrhythmogenic Cardiomyopathy with Multi-Ion Channel Remodeling and Ankyrin-G Scaffold Disruption"

Supplementary Tables 1 to 3.

Supplementary Figures 1 to 4.

**Supplementary Tables**

**Supplementary Table 1. RT-qPCR primers for gene expression**

| **Gene** | **Species** | **NCBI ID** | **NCBI accession number** | **Primer sequence**  **(5’ to 3’)** | **Product size (bp)** |
| --- | --- | --- | --- | --- | --- |
| **β-Actin** | Rat | 81822 | NM_031144 | F: TGTCACCAACTGGGACGATA | 165 |
|  |  |  |  | R: GGGGTGTTGAAGGTCTCAAA’ |  |
| **BVES** | Rat | 365603 | NM_001077590 | F: ACTCGGTGTTCTTGGGTAT | 232 |
|  |  |  |  | R: GCTAAGACGGTCATCAACT |  |
| **ANKG** | Rat | 361833 | NM_001033984 | F: ACACAGCCTTACACATCGCATCC | 147 |
|  |  |  |  | R: TGACGACTTCCAGGTGGTTCTCC |  |

**Supplementary Table 2. QTc interval calculations using multiple formulas**

| **Formula** | **Formula definition** | **Normal sinus (HR 70–72 bpm)** | | **Bradycardia (HR 43–50 bpm)** | |
| --- | --- | --- | --- | --- | --- |
|  |  | **Absolute QT** | **Calculated QTc** | **Absolute QT** | **Calculated QTc** |
| **Bazett^1^** | QT / √RR | 360 – 380 ms | 389 – 416 ms | 360 – 380 ms | **305 – 347 ms** |
| **Fridericia^2^** | QT / ∛RR | 360 – 380 ms | 379 – 404 ms | 360 – 380 ms | **322 – 358 ms** |
| **Framingham^3^** | QT + 154(1-RR) | 360 – 380 ms | 382 – 406 ms | 360 – 380 ms | **299 – 349 ms** |
| **Hodges^4^** | QT + 1.75(HR-60) | 360 – 380 ms | 378 – 401 ms | 360 – 380 ms | **330** **– 363 ms** |

***Note:*** *Calculations are based on the proband’s clinical ECG recordings. While QTc intervals remained normal at normal sinus heart rates, all four correction methods demonstrated a paradoxical shortening into the diagnostic Short-QT range when presented with bradycardia.*

**Supplementary Table 3. Rare POPDC1 Variants Identified in Early-Onset AF and DCM Cohorts**

| Patient ID |  | Variant (c. / p.) | Type | Zygosity | MAF (EAS)^†^ | Key Clinical Features |
| --- | --- | --- | --- | --- | --- | --- |
| Early-onset AF cohort (n=222) | | | | | | |
| Patient AF-01 |  | c.1009G>T (p.Glu337*) | Nonsense | Heterozygous | 0.0000 | Persistent AF |
| Patient AF-02 |  | c.56C>A (p.Pro19His) | Missense | Heterozygous | 1.6 × 10^-5^ | Prominent J-wave elevation; long-standing persistent AF; Recurrent AF recurrence after radiofrequency ablation |
| DCM cohort (n=344) | | | | | | |
| Patient DCM-01 |  | c.642A>C (p.Lys214Asn) | Missense | Heterozygous | 0.0000 | Heart failure symptoms and PVC |
| Patient DCM-02 |  | c.313A>C (p.Ile105Leu) | Missense | Heterozygous | 2.2 × 10^-5^ | SCD at age 50 (VF documented); Severe LV dilation and left ventricular noncompaction |

Abbreviations: AF, Atrial Fibrillation; DCM, Dilated Cardiomyopathy; SCD, Sudden Cardiac Death; VF, Ventricular Fibrillation; MAF, Minor Allele Frequency; EAS, East Asian Population; PVC, premature ventricular contraction

† MAF data derived from GnomAD v4.1.0 (East Asian).

**Supplementary Figures**

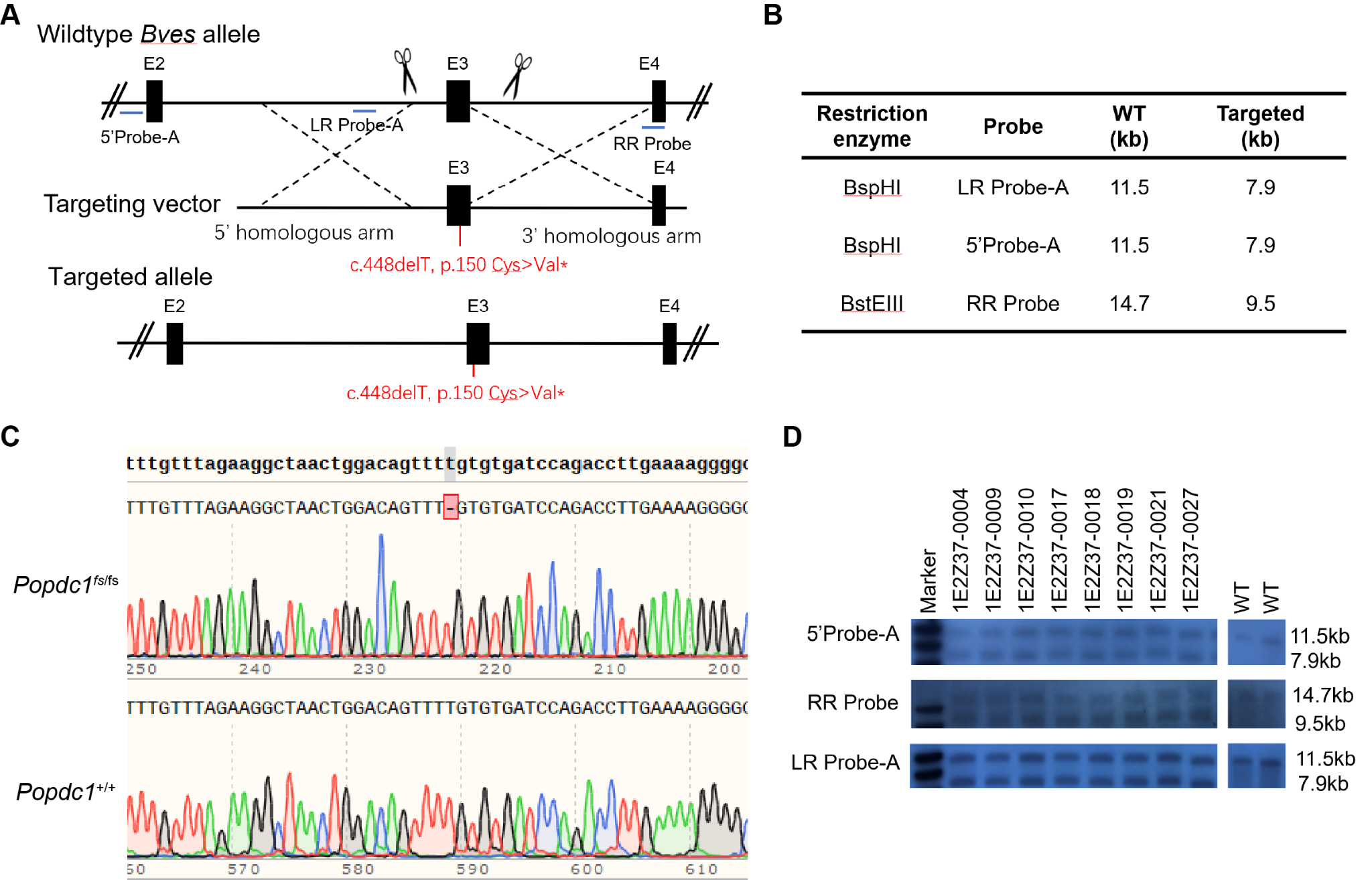

**Supplementary Figure 1. Generation and structural characterization of the orthologous *Popdc1*^fs/fs^ rat model.** (A) Schematic of the CRISPR/Cas9 targeting strategy used to introduce the c.448delT mutation. (B) Restriction enzymes of the WT Bves locus and targeted locus. (C) Sanger sequencing validation confirming the frameshift genotype in Popdc1fs/fs mutants. (D) Southern blot validation of genotype sequence compared to WT.

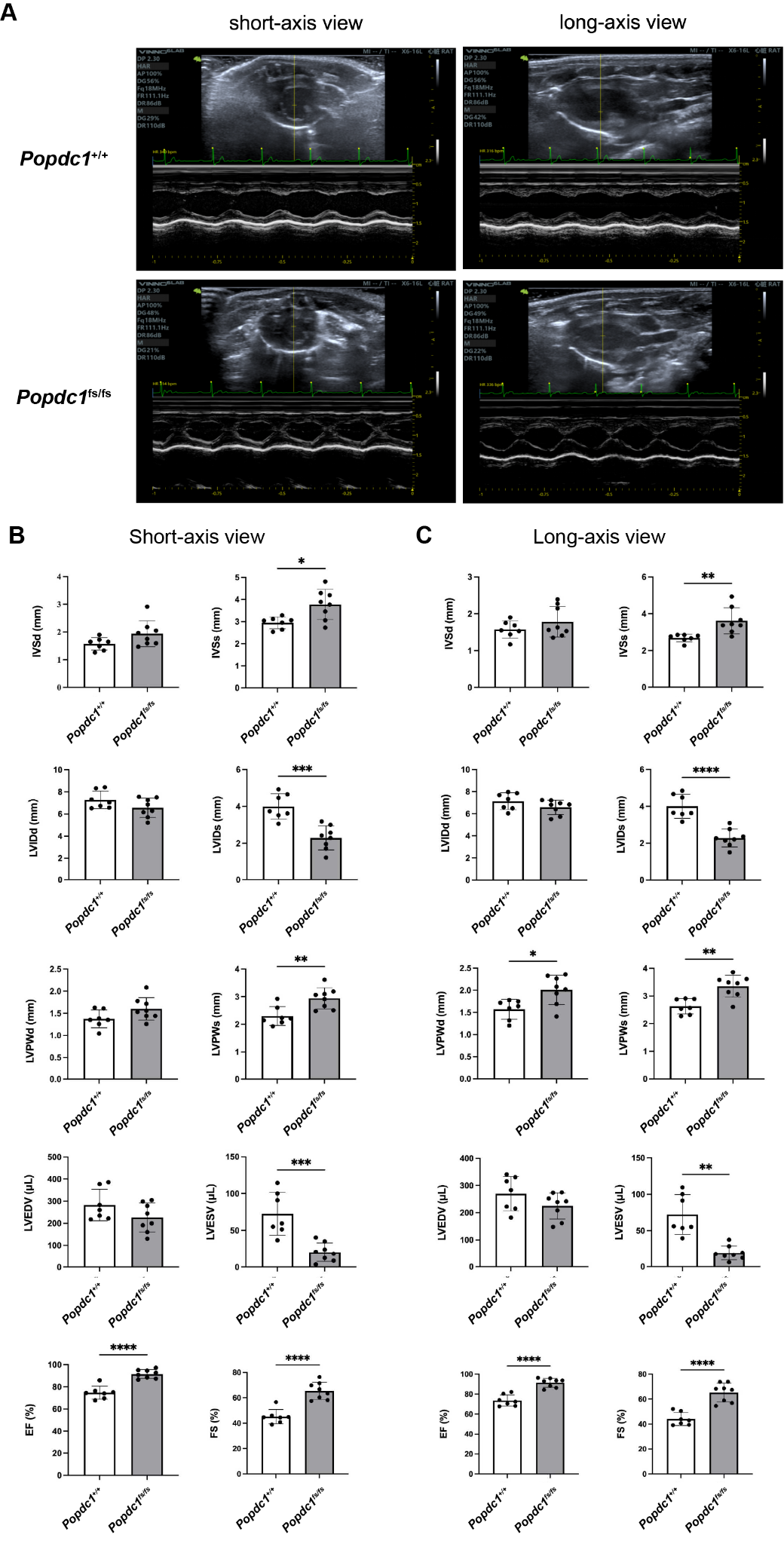

**Supplementary Figure 2. *Popdc1*^fs/fs^ rats demonstrate hypercontractility and LV hypertrophy phenotype evident during systole.** (A) Representative M-mode images showing the development of LV wall thickening (hypertrophy-like pattern) in mutants compared to WT. (B-C) Cardiac phenotype during short-axis view (B) and long-axis view (C) from ultrasound images (n=7-8/group). Definitions are as follows: intra-ventricular septum, IVS; LV inner diameter, LVID; LV posterior wall, LVPW; LV end-diastolic volume, LVEDV; LV end-systolic volume, LVESV; measured during diastole, d and systole, s. Data expressed as mean ± SD. Statistical analyses by unpaired Student’s t-test (B, C). *P<0.05, **P<0.01, ***P<0.001 respectively.

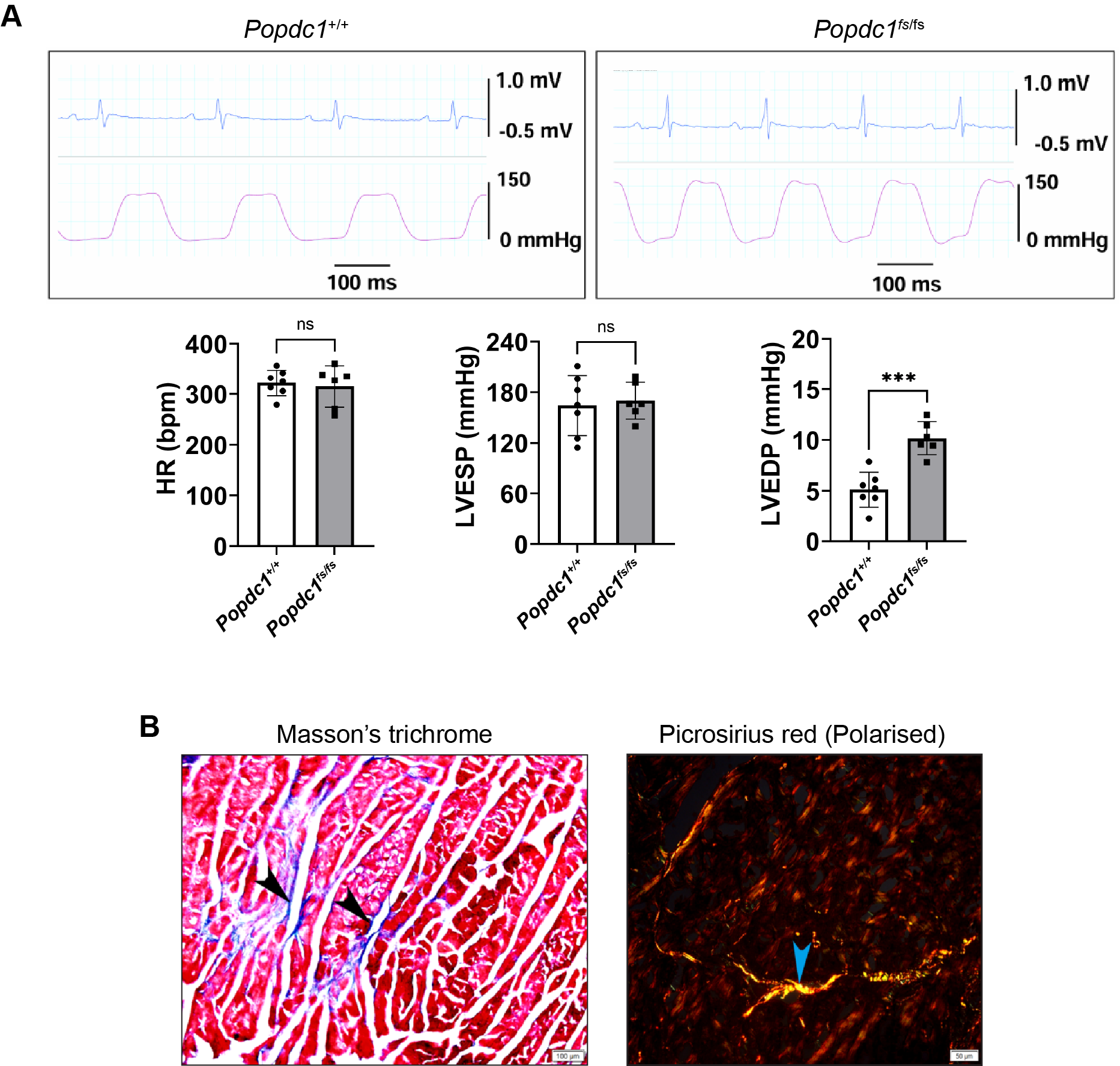

**Supplementary Figure 3. *Popdc1*^fs/fs^ rats demonstrate impaired relaxation and diffused interstitial fibrosis.** (A) Representative ECG (top) and concurrent LV pressure traces (bottom) demonstrated elevated LV end-diastolic pressure (LVEDP) but unchanged LV end-systolic pressure (LVESP) and heart rate. (B) Representative Masson’s trichrome and picrosirius red staining for collagen content reveals prominent interstitial fibrosis in *Popdc1*^fs/fs^ hearts (n=3). Scale bar: 100 µm. Data are mean ± SD. Statistical analyses by unpaired Student’s t-test.

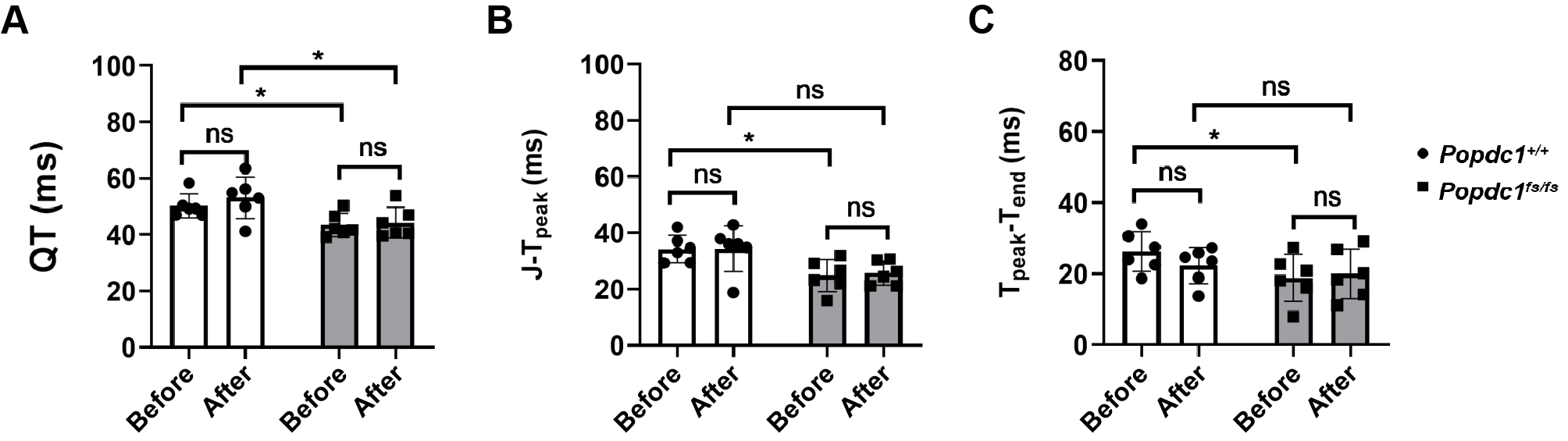

**Supplementary Figure 4. Chronotropic incompetence and blunted adrenergic sensitivity due to swimming in *Popdc1*^fs/fs^ rats.** (A-C) ECG indices demonstrating reduced QT interval (A), J-Tpeak interval (B), and Tpeak-end (C) in *Popdc1*^fs/fs^ rats before swimming exercise. Additionally, the short QT interval remained refractory to swimming in *Popdc1*^fs/fs^ rats . Data are mean ± SD. Statistical analyses by paired t-test. *P<0.05, ns = not significant.
